# M1A within cytoplasmic mRNAs at single nucleotide resolution: A reconciled transcriptome-wide map

**DOI:** 10.1101/308437

**Authors:** Schraga Schwartz

## Abstract

Following synthesis, RNA can be modified with over 100 chemically distinct modifications, and in recent years it was shown that processing, localization, stability and translation of mRNAs can be impacted by an increasing number of these modifications. A modification that recently gained attention is N1-methyladenosine (m1A), which is present across all three domains of life. Recently, two studies - one of them ours - developed conceptually similar approaches to map m1A in a transcriptome-wide manner and at single nucleotide resolution. Surprisingly, the two studies diverged quite substantially in terms of their estimates of the abundance, whereabouts and stoichiometry of this modification within internal sites in cytosolic mRNAs: One study reported it to be a very rare modification, present at very low stoichiometries, and invariably catalyzed by TRMT6/61A. The other found it to be present at >470 sites, in dozens of which at relatively high levels, and in the vast majority of cases these sites were highly unlikely to be substrates of TRMT6/61A, suggesting that additional methyltransferases are active on cytosolic mRNAs. Here we aim to reconcile the contradictions between the two studies, primarily by reanalyzing and re-annotating the set of sites identified in the latter study. We find that the vast majority of sites detected in this study originate from duplications, misannotations, mismapping, SNPs, sequencing errors, and a set of sites originating from the very first transcribed base (‘TSS sites’). We raise concerns as to whether the TSS sites truly reflect m1A originating from the first transcribed base. We find that only 53 of the sites detected in this study likely reflect bona-fide internal modifications of cytoplasmically encoded mRNA molecules. The vast majority of these are likely to be TRMT6/TRMT61A substrates, and are typically modified at low to undetectable levels. We conclude that within cytosolic mRNAs, m1A is a rare internal modification where it is typically catalyzed at ultra-low stoichiometries via TRMT6/TRMT61A. Our findings offer a clear and consistent view on the abundance and whereabouts of this modification, and lays out key directions for future studies in the field.

## Introduction

Post-transcriptional modifications of RNA form an emerging layer of regulation of gene expression, analogous in potential importance to post-translational modifications of proteins. Over 100 modifications exist throughout the three domains of life. While these modifications were traditionally studied in the highly abundant - and hence biochemically tractable - tRNA and rRNA molecules, in recent years high-throughput sequencing approaches have allowed to generate transcriptome-wide maps of RNA modifications. These maps have revealed that some modifications are widespread also in other classes of RNA, most notably in mRNA (Carlile et al., 2014; Dominissini et al., 2012; Li et al., 2015; Meyer et al., 2012; Schwartz et al., 2013, 2014a), where they can impact mRNA processing, localization, stability and translational efficiency (Haussmann et al., 2016; Lence et al., 2016; Meyer et al., 2015; Schwartz et al., 2014b; Shi et al., 2017; Wang et al., 2014, 2015; Zhou et al., 2015).

Nearly two years ago, two reports, coupling the use of an anti-m1A antibody with RNA-sequencing, collectively reported the identification of >7000 putative m1A harboring regions in one study (Dominissini et al., 2016) and nearly 1000 regions in another (Li et al., 2016). Both studies found m1A to originate primarily from 5’ UTRs, and particularly near start codons or first exon-exon junctions (Dominissini et al., 2016; Li et al., 2016). One study found the putative m1A sites to be associated within a degenerate GC rich consensus (Dominissini et al., 2016); the other observed an enrichment for a degenerate purine rich motif (Li et al., 2016). Both studies lacked the ability to detect m1A at single nucleotide resolution, and did not identify enzymes catalyzing formation of m1A on mRNA, and hence could not directly validate these sites, nor assay their functions and mechanisms of action.

Recently, two studies - one of which by our group - developed conceptually similar approaches for mapping m1A at single nucleotide resolution (Li et al., 2017; Safra et al., 2017). Rather than relying on enrichment of m1A-containing fragments upon m1A-IP, these two studies relied on misincorporation patterns introduced upon reverse transcription of m1A containing RNA (Hauenschild et al., 2015). The studies used additional measures to ensure that the misincorporation patterns were specific. Both studies measured misincorporation levels in the input RNA, and upon pulldown using the anti-m1A antibody, and sought misincorporation signals enriched in the latter. To achieve additional stringency, both studies further measured misincorporation levels in an immunoprecipitated sample subjected to a treatment that eliminates, or reduces, m1A levels: In one case the samples were subjected to Dimroth rearrangement, a chemical treatment that converts m1A residues into m6A residues and thereby reduces misincorporation levels (Safra et al., 2017). In the other case, elimination of m1A was achieved via the employment and careful calibration of an RNA demethylating enzyme, leading to almost complete elimination of m1A levels (Li et al., 2017). The Safra et al. study further developed an approach relying on reverse transcription using an enzyme that predominantly leads to premature truncation of reverse transcription, as an additional control (Safra et al., 2017).

Reassuringly, the two studies converged on some of their findings. Both identified sites in the cytoplasm, sharing an identical sequence and structural features, and found them to be modified by the TRMT6/TRMT61A complex at an identical sequence and structural motif as found in position 58 of tRNA, the well-characterized substrate of this complex. Both studies further found that m1A was present at a number of sites within mitochondria, leading both of them to focus in particular on the same site in ND5, a mitochondrially encoded gene forming part of complex I of the respiratory complex. Furthermore, both studies found that m1A within internal positions of mRNA represses translation.

Nonetheless, the two studies diverged substantially in terms of their estimates of the abundance, whereabouts and stoichiometry of m1A. Safra et al. reported the identification of eight m1A sites in cytosolic mRNAs and lncRNAs, most of which were estimated to be modified at very low levels (most were undetectable in the Input samples), and all of them catalyzed via TRMT6/TRM61A. In addition, this study reported that m1A-IP enriches for the 5’ end of genes, but that this enrichment does not originate from 5’ UTRs, or from the start codon, or from the first splice junction as previously reported (Dominissini et al., 2016; Li et al., 2016), but rather from the very first transcribed base (‘TSS sites’). Finally, this study did not detect evidence for m1A presence at the TSS sites on the basis of analysis of misincorporation patterns, and hence left open the question as to whether these sites originate from m1A (or an m1A-derivative) at the TSS or from antibody promiscuity. In contrast, Li et al. reported a total of 474 sites, of which only 53 harbored a TRMT6/61A motif. They further classified a total of 277 sites as originating from the 5’ UTR, only 24 of which mapping to the first transcribed nucleotide. The findings of Li et al. suggest that (1) m1A at internal positions on mRNA is substantially more prevalent than reported by the Safra study, (2) TRMT6/61A only has a minor role in shaping the m1A landscape, suggesting that other enzymes remain to be discovered, (3) m1A at internal sites can be present at considerable stoichiometry: 76 of the mRNA sites reported by Li et al. have mean misincorporation rates >20% within the Input - unenriched - fractions, and the authors further emphasize that misincorporation rates likely substantially underestimate the true m1A levels. The findings of Li et al thus provide support to the two original publications on m1A, which characterized this modification to be widespread at internal sites within mRNAs, primarily within 5’ UTRs, and to be present at a stoichiometry of ~20% (Dominissini et al., 2016; Li et al., 2016).

Here, we seek to reconcile the points of divergence between the two studies, primarily by reanalyzing and re-annotating the set of sites identified by Li et al. We find that 53 of the 474 sites are likely to reflect bona-fide internal modifications of cytoplasmically encoded mRNA molecules. All of these sites harbor TRMT6/TRMT61A consensus motifs and are modified at low to undetectable levels, in the absence of IP. The remaining sites correspond to (1) sites appearing redundantly within this dataset, (2) sites originating from tRNA or mitochondrial RNA which were mismapped or misannotated as mRNA, (3) Genomic SNPs or sequencing errors, or (4) sites originating from the very first transcribed base (‘TSS sites’). We raise concerns as to whether the TSS sites truly reflect m1A modification of the first transcribed base, but a definitive answer as to the nature of the signal at the TSS awaits interrogation using biochemical approaches.

## Results

### Reclassification of putative m1A sites

Li et al aligned the sequencing reads to a transcriptomic reference, rather than to the human genome, and identified 474 putative m1A sites. 53 of these sites were classified as putative TRMT6/61A substrates, 24 as originating from the first transcribed nucleotide, and the remaining site were unclassified (**Fig. 1A, left panel**). To allow comparison of these sites to other datasets, we first converted the transcriptomic coordinates to genomic coordinates. In this context we eliminated from this dataset 3 sites which did not form part of the current Refseq database and 37 sites that did not have an ‘A’ at the reported site in this annotation (these discrepancies likely reflect updates of the Refseq database; The Li et al study does not specify the precise Refseq version that was used). Importantly, given that 37 of the 474 sites did not harbor an A at the reported position, we estimate a minimal false conversion rate of ~7.8%, and thus estimate that further ~39 of the remaining sites analyzed here do not reflect the original sites identified by Li et al. We then re-classified and filtered the 434 remaining sites (**Fig. 1A, right panel**), as follows:

- **Redundant entries**: 82 of the remaining sites were redundant entries, i.e. entries appearing at least twice in the reported dataset (in one extreme case the same site appears 8 times in the table). Although Li et al eliminated transcripts harboring identical Refseq IDs, this was not sufficient to eliminate such redundancies, as transcripts with distinct refseq IDs can also overlap the same genomic locus, hence resulting in an artificial inflation of the number of detected sites.
- **Mitochondrial pseudogenes and tRNAs**: 22 of the remaining sites mapped to mitochondrial pseudogenes, also known as nuclear mitochondrial DNA segments (NUMTS) and 5 sites originated from tRNAs in regions which had overlapping annotations also with mRNAs. In both cases, the identified sites likely arise from the mitochondrial RNA and from tRNAs (both of which have well documented m1A sites), respectively, and not from the mRNA annotated to overlap them. Indeed, visual inspection of the alignment patterns at these sites further confirmed that the mapping patterns within these genes were inconsistent with the annotation of the mRNAs annotated at this site, and hence likely stem from reads originating from the mitochondria, that underwent mismapping to the NUMTs, and reads originating from tRNAs that were misannotated as mRNAs.
- **TRMT6/TRMT61A substrates**: Li et al. required a strictly defined consensus sequence (GTTCRA) in order to classify a site as TRMT6/61A dependent, with no structural constraints. As we previously reported, TRMT6/61A substrates require both a consensus sequence and a structure, although there is considerable flexibility with regard to both (Safra et al., 2017). We thus classified sites as harboring a TRMT6/TRM61A motif if they harbored both a slightly relaxed motif (G[ATC]TCNA) and a stem of at least three bp, including basepairing between position −5 and +3 with respect to the m1A site, −6 and +4; These two basepairings are the most critical for the formation of m1A (Safra et al., 2017). Sites with only one of these two criteria were still considered TRMT6/TRMT61A substrates, unless they were within the first 200 bp in which case they were annotated as TSS sites (see below). 53 sites matched these criteria (note that although this number is identical to the one reported by Li et al, the catalog of sites is distinct and both includes sites that were not originally included by Li et al due to our relaxed consensus requirements, and eliminates sites that were reported by Li et al but which originate from tRNAs). Of note, 35 of these sites (66%) formed part of the dataset of 495 sites that we reported to acquire m1A upon overexpression of TRMT6/TRMT61A (Safra et al., 2017), lending strong evidence to the notion that these sites are highly enriched in TRMT6/TRMT61A substrates.
- **TSS sites**: Li et al identified many sites mapping to 5’ UTRs, but classified a site as originating from the first transcribed nucleotide only if it mapped to the very first base of the *annotated* transcript. This approach will result in a substantial underestimate of the number of sites originating from the first transcribed base, as transcription typically does not begin at a single nucleotide but across a region comprising several dozens of nucleotides, of which the refseq annotation chooses a representative site (Carninci et al., 2006; Plessy et al., 2010; Takahashi et al., 2012). Given that the library preparation procedure employed by the Li et al study allows capturing the first transcribed base at single nucleotide precision, we first inspected the reads at the identified sites. This analysis revealed that the mutations, when present in the sites annotated as 5’ UTR, almost invariably occurred at the first base of the read (**Fig. 1B**), strongly suggesting that they occur at the first transcribed nucleotide. To perform this analysis in a systematic manner, we examined CAGE data, a technique dedicated to obtaining single nucleotide resolution mapping of TSS. ENCODE CAGE data from A549 cells (ENCODE CAGE data for HEK293 does not exist) reveals that 79% of the non-TRMT6/61A sites which occur within 200 bp of the annotated start site have at least 1 CAGE tag associated with them, indicating that they serve at least occasionally as the first transcribed nucleotide (in contrast, only 3% of sites classified as TRMT6/61A sites had one such tag). We thus classified 196 sites residing within the first 200 bp as putative TSS sites.
- **Other sites**: 76 (of the 434) sites do not readily fall into any of the above defined criteria. 43 of these sites had misincorporation ratios in the Input samples exceeding 10%, and we individually inspected each of them. We found only 2 cases of convincing misincorporation patterns, consistent with m1A (i.e. high in IP, lower in Input, low upon demethylase treatment). Both sites are with very high likelihood TRMT6/61A substrates, but have minor variations in the motif/structure causing them not to be classified as such. Of the remaining sites:

○ 11 originate from the first transcribed nucleotide (with misincorporations present only at the first base of the read) in a manner inconsistent with the Refseq annotation;
○ 4 sites have a known SNP at the modified sites, and the identified misincorporations merely reflect the presence of two alleles;
○ 6 sites occurred within poly(C) stretches, and the misincorporation pattern in them were always A->C, which is highly atypical of the m1A signature. These sites also generally exhibited very poor enrichment upon m1A-IP. Moreover, mutations to C are in some of the cases also observed in adjacent sites which do not harbor an A, but are also embedded within poly(C) stretches. These sites thus likely reflect Illumina errors at homopolymeric C runs, where the basecaller appears to call the homopolymer base also in the cycle following the homopolymer. Such a phenomenon has been previously attributed to limited handling of (pre-) phasing in homopolymeric stretches and to signals accumulating over cycles due to incomplete cleavage of fluorophore (Chang et al., 2014; Ledergerber and Dessimoz, 2011; Whiteford et al., 2009).
○ 3 sites had ‘noisy’ alignment patterns, i.e. there were many ‘misincorporations’ either in the reads supporting them or in the adjacent region. In 2 of these cases the sites are within an Alu repeat, and hence the noisy alignments likely reflect reads originating from elsewhere.
○ 17 sites either harbored no reads or no misincorporations when aligned to the human genome. The discrepancy between these patterns and the reported ones are likely a result of differences between our annotation and the one used by Li et al, resulting the above-described ‘false conversion rate’. It is furthermore possible that misincorporations reported at some of these sites are due to reads that were mapped to a region in the transcriptome, despite originating from a different region in the genome. This is a key limitation when mapping reads to the transcriptome, as it is heavily dependent on the annotation and can force alignments even in the presence of better genomic alignments.

**Figure 1.**
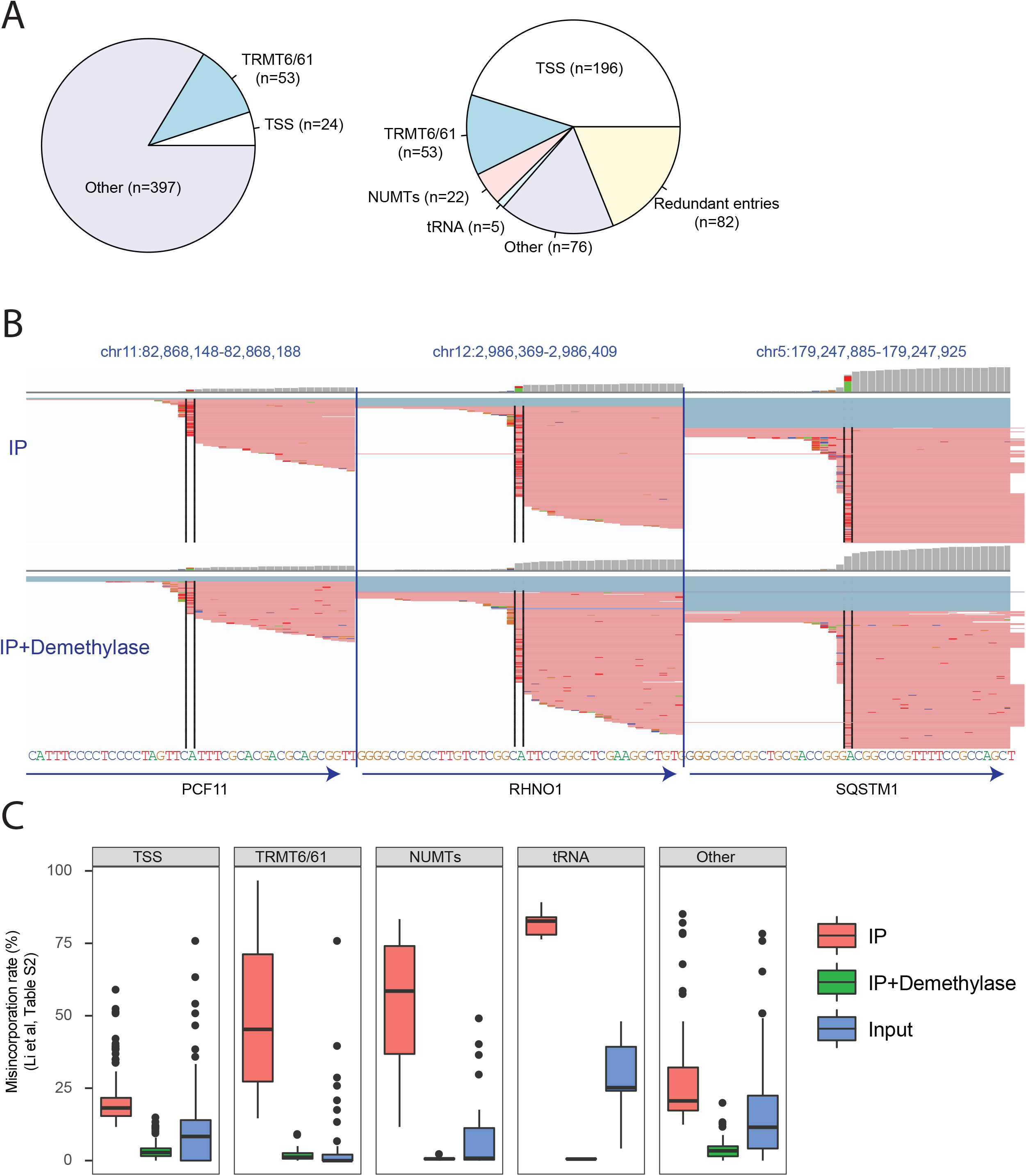
Re-annotation and characterization of putative m1A sites identified by Li et al. (**A**) Pie charts depicting the original classification of the putative m1A sites (left) and the re-classified sites (right). (**B**) Genome browser snapshots of three genomic loci, harboring putative m1A peaks originally classified by Li et al as originating from the 5’ UTR. The genomic reference is depicted at the bottom. The red lines overlapping each locus are reads, with bases diverging from the reference encoded by different colors (green - A, blue - C, orange - G, red - T). The black bars enclose the putative m1A sites. Although the gene annotation (depicted by arrows on the bottom) all indicate that the ‘official’ gene start site is upstream of the modified site, it is apparent that all the misincorporations occur at the first base of the sequencing read. The top panel presents the mismatches upon m1A-IP, the bottom upon IP and demethylase treatment. (**C**) Distribution of misincorporation rates across Input, IP, and IP + Demethylase treated samples across the five different classes into which the Li et al sites were re-classified, on the basis of the misincorporation ratios reported in Table S2. These quantifications highlight strong enrichment of TRMT6/61A sites, but much reduced enrichment of TSS sites.

The re-annotated and reclassified table of 352 non-redundant sites is available as **Table S1**.

### TRMT6/TRMT61A sites are typically modified at very low levels

We next examined the misincorporation profile at the identified TRMT6/TRMT61A sites in Input, IP, and IP + Demethylated samples. For each site we refer to the mean misincorporation ratio, based on two experimental replicates reported in Supplementary Table S2 in the manuscript by Li et al. This analysis reveals that the median misincorporation rate for TRMT6/TRMT61A sites in the Input samples was 0% (but 45% after IP); Only 8 sites had misincorporation rates exceeding 5% (**Fig. 1C**). Moreover, inspection of each of these 8 sites revealed only a single site, in the PRUNE gene 5’ UTR, that had convincing misincorporation rates (16%) in the Input sample; Note that this is also the one cytoplasmic site identified and followed up on in the Safra et al manuscript, due to the higher misincorporation levels detected there as well (Safra et al., 2017). In all other cases, upon genomic mapping of the Li et al. reads in the input samples we either observed very low read counts at these sites, or no reads at all. The paucity or lack of coverage is likely due to false conversion rates or to issues with alignment to transcriptomes instead of genomes, as above, and in few cases may also be due to the lower depth to which the Input samples were sequenced. While it thus remains possible that some of them are modified to a low extent, which could be quantifiable via targeted sequencing, overall the data does not provide evidence for m1A being present at substantial levels.

Are misincorporation rates an accurate proxy of m1A stoichiometries? Given that misincorporation rates at m1A are sequence dependent (Hauenschild et al., 2015), we assessed the *maximal* incorporation rate that can be expected on the basis of the TRMT6/TRMT61A consensus sequence. For this we utilized the misincorporation rates within tRNAs at position 58 - which harbor the identical consensus sequence as the positions identified in mRNA, obtained from Supplementary Table 1 released by Li et al. We found that the median misincorporation rates at this site upon m1A-IP was 83%. Even under the conservative assumption that the IP was 100% efficient, and that it isolated only m1A containing molecules, this would imply that misincorporation rates provide a mild underestimate of m1A stoichiometry, at the order of ~20% of the methylation levels.

### Misincorporations at the TSS - m1A or not m1A?

Although the m1A antibody clearly enriches for TSS as supported by all studies applying it (Dominissini et al., 2016; Li et al., 2016, 2017; Safra et al., 2017), several considerations raise important concerns as to whether the TSS sites truly harbor m1A, or whether the enrichment may be attributed to an m1A derivative or potentially also to promiscuous binding by the antibody: (**1**) although the TSS sites have substantially higher levels of misincorporations in the Input samples (median: 8.3%), the sites reported by Li et al are only poorly enriched upon IP (median: 18%), in stark contrast to TRMT6/61A substrates (**Fig. 1C**). (**2**) Bona fide m1A sites are expected to lead to premature truncation of reverse transcription when reverse transcribed using SuperScript; The rates of such truncations are expected to be decreased upon Dimroth rearrangement which converts m1A to m6A, which no longer induces the stop (Safra et al., 2017). Using our published m1A-seq-SuperScript dataset, we found no significant change in stop rates at the detected sites between samples subjected to IP and samples subjected to IP + Dimroth treatment (Paired T test, P=0.12) (**Fig. 2A, left panel**); In contrast, there was a highly significant drop in stop rates when performing this analysis for sites in the TRMT6/61A group, as a control (Paired T test, P=0.006) (**Fig. 2A, right panel**). (**3**) Similarly, when quantifying misincorporation rates in the m1A-Seq-TGIRT dataset generated in Safra et al at the set of putative sites identified in Li et al, we do not find evidence for increased misincorporation rates at the TSS sites (**Fig. 2B, left panel**), in stark contrast to the TRMT6/61A sites (**Fig. 2B, right panel**). (**4**) There is a very poor overlap between the set of TSS sites reported here and the set of m1A peaks identified by the two original studies. Of the 901 ‘m1A peaks’ reported by the authors in their previous study (Li et al., 2016), only 7 overlap with the TSS peaks; And of 2129 ‘m1A peaks’ reported in Hek293 cells (Dominissini et al., 2016), only 13 overlap with the TSS sites. Thus, for the overwhelming majority of the previously reported peak regions that were enriched by the anti-m1A antibody, no evidence of misincorporation is observed. Given that the methodology developed by Li et al is highly sensitive, and - as exemplified by the TRMT6/61A substrates - is able to detect sites that are modified at levels close to 0%, and given that the peak regions typically have extremely deep coverage by virtue of being enriched upon m1A-IP, the lack of detected misincorporations across the vast majority of these sites thus strongly indicates that such misincorporations typically do not occur at these sites.

**Figure 2.**
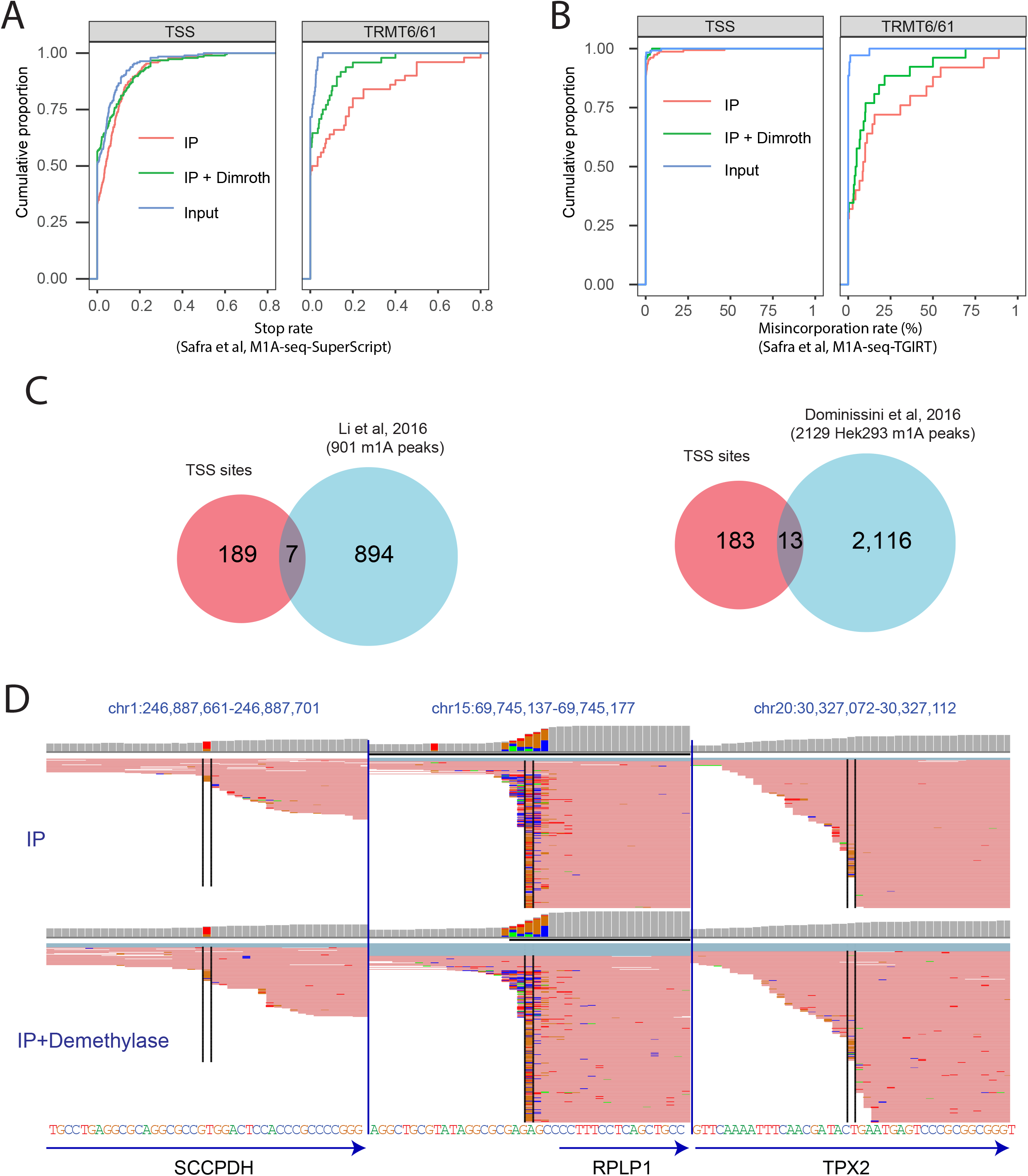
Characterization of TSS sites. (**A**) Analysis of the ‘stop rate’, defined as the fraction of reads beginning at a site (indicating reverse transcriptase termination) divided by the number of reads overlapping it, on the basis of the m1A-seq-SuperScript dataset released by (Safra et al., 2017). SuperScript leads to a high termination rate of reverse transcription (and lower misincorporation rate) and hence serves as an orthogonal methodology for detecting m1A. The rate of termination does not differ significantly between IP and IP+Dimroth across TSS sites (left panel) in stark contrast to TRMT6/61A sites (right panel) used as positive control. (**B**) Analysis of the misincorporation rate at the Li et al sites on the basis of the Safra et al dataset. The misincorporation rate does not differ between the samples regarding TSS sites (left panel) in contrast to the TRMT6/61A sites (right panel). (**C**) Venn diagram depicting the overlap between the TSS sites, as classified here, and the sets of ‘m1A peaks’ identified as enriched in (Li et al., 2016) (left panel) and (Dominissini et al., 2016) (right panel) respectively. (**D**) Genome browser snapshots (as in **Fig. 1B**) of three arbitrarily selected regions around the transcription start sites of genes, in which misincorporations enriched upon IP with respect to IP + Dimroth were **not** identified. In all cases, misincorporations can be viewed both at the position marked by the black lines, but often also in nearby positions. In the left and right panels the highlighted position does not contain an A. In all cases, the mismatches with respect to the genome occur at the very first base of the read (in some cases they encompass the first 2 or 3 bases of the read). The misincorporations are present both in IP and IP+Dimroth treated samples.

One potential source for the observed misincorporations at the very first position is the potential for non-templated nucleotide incorporations in the context of reverse transcription: It is a well characterized phenomenon that reverse transcriptases can include such non-templated additions (Chen and Patton, 2001). Indeed, inspection of the misincorporations patterns across many TSS, both enriched and not enriched via the anti-m1A antibody, reveals abundant ‘misincorporations’ occuring at the very first position (**Fig. 2D**); Moreover, such misincorporations are not exclusive to first bases harboring an ‘A’ (**Fig. 2D, middle panel**) but also occur at transcripts beginning with other bases (**Fig. 2D, left and right panel**). While these misincorporations are typically not enriched upon pulldown nor eliminated upon treatment with the demethylase, by explicitly filtering for such sites from across the entire genome, such sites may potentially be spuriously detected. We should further note that due to these widespread misincorporations at the very first position, in the Safra et al manuscript we made use of the default alignment option of the STAR aligner (Dobin et al., 2013), which utilizes soft clipping: This option trims bases from their beginning (and ends), to maximize the alignment score. Therefore, misincorporations occuring at the very first base of the read are soft clipped, and hence not taken into account in the context of misincorporation detection.

### High overlap in detection of TRMT6/61A sites - but not of TSS sites - between computational approaches

Why did the Li et al approach result in identification of 53 TRMT6/61 substrates, and the Safra et al approach only in 8? A trivial explanation would be that this is a consequence of the increased coverage in the Li et al study. Due to increased sequencing depth, increased read length and improved depletion of rRNA sequences, the median coverage per gene in Li et al was 10.2 fold higher in the two IP samples in comparison to the Safra et al dataset, and such enrichment should allow detection of sites in more poorly expressed genes. However, the differences in the computational approaches employed by the two groups could also lead to differential detection. To directly assess whether the approach employed by Safra et al has reduced sensitivity in detecting m1A sites, we applied the computational approach developed in the Safra et al manuscript to the raw data obtained from the Li et al studies. This approach uncovered a total of 65 sites within mRNAs. Of these, 51 (78%) underwent classification as TRMT6/61A sites on the basis of the above-defined criteria; Visual examination of the remaining ones revealed that at least 8 of them are likely to be TRMT6/61A candidates as well, but with atypical secondary structures causing them to undergo misclassification. Among the putative TRMT6/61A substrates, 32/53 (~60%) of the TRMT6/61A sites identified by Li et al also identified by the Safra et al approach. Of note, only 5 of the 196 (2.5%) sites within the TSS class were identified using the Safra et al approach, and inspection of all of these revealed that they are all likely to be TRMT6/61A substrates (4 of these 5 sites formed part of the sites previously identified by Safra et al upon TRMT6/61A overexpression). Thus, overall the computational approaches by Safra et al and Li et al yield a similar number of putative TRMT6/61A substrates when applied to the same dataset, and with a reasonable extent of overlap between them. In stark contrast, despite the fact that ~4 fold more sites in the Li et al dataset are classified as TSS sites than TRMT6/61A sites, essentially none of these sites is detected when applying the Safra et al computational approach.

## Conclusion

Re-analysis of the Li et al data thus reveals a view highly similar to the one identified in Safra et al: Both datasets demonstrate that within internal positions in cytosolic mRNA molecules, m1A is nearly exclusively found within a typical sequence and secondary structure, where it is catalyzed by TRMT6/TRMT61A. Thus, there does not seem to be evidence in support of an additional m1A methyltransferase, acting on a distinct subset of internal sites on mRNA. While the higher coverage in the Li et al facilitated the identification of more sites compared to the Safra et al study, the characteristics of these sites in terms of their sequence/structural requirements, their stoichiometries, and the enzymatic complex catalyzing their formation are essentially identical.

It is likely that, in immunoprecipitated samples sequenced to even greater depth, it will be possible to find evidence for the existence of even more TRMT6/61A sites; In fact, it is possible that it will be possible to find evidence for m1A at very minor levels for all ~400 sites identified in the Safra et al study upon overexpression of TRMT6/61A. An interesting analogy is to A->I editing, where - based on ultradeep sequencing of selected loci - it was estimated that over 100 million editing residues may exist in the human genome, albeit the vast majority at levels <<1% (Bazak et al., 2014). The fact that m1A levels within mRNA TRMT6/61A sites are - in the overwhelming majority of cases - present at very low stoichiometries suggests that m1A at internal positions is unlikely to play a major role in regulation of gene expression. This, however, does not preclude a potentially more dramatic role played by this modification in mitochondrial mRNAs, where this modification is found, at least in one case, at a higher stoichiometry (Li et al., 2017; Safra et al., 2017).

An important, future direction to better understand is the nature of the signal at the transcription start site. Our reanalysis of the Li et al dataset strongly suggests that the enrichment previously reported to originate from the 5’ UTR, or from the first exon-exon junction (Dominissini et al., 2016; Li et al., 2016, 2017) in fact originates from the first transcribed base, consistent with our previous report (Safra et al., 2017). The low-level enrichment observed for TSS sites, the absence of RT-induced truncations, and the fact that such sites can only be detected at a very low fraction of the previously reported m1A-peaks raise concerns as to whether these sites truly reflect m1A occuring at the first transcribed nucleotide. These concerns are compounded by the fact that detection of these sites relies on mismatches occuring at the first position of the read which is particularly prone to non-templated nucleotide incorporations occuring as part of reverse transcription. This notwithstanding, none of these observations provide an answer as to why certain TSSs are enriched with the anti-m1A antibody, nor do they definitely rule out that m1A - or an m1A derivative - is present at the TSS. The reports that presence of such enrichment - whatever its nature - correlates positively with translation (Dominissini et al., 2016; Li et al., 2017) are intriguing. We anticipate that a definitive response to these questions will arise from biochemical approaches.

## Methods

The re-analysis of this study primarily relies on Table S2 in the Li et al study. Transcriptomic coordinates were mapped to genomic coordinates using an in-house script, on the basis of an updated Refseq annotation (downloaded on Dec. 12, 2017 from the UCSC Table browser). The ‘tRNA Genes’, ‘NumtS’ and ‘Common SNPs (147)’ tables providing the coordinates of tRNAs, mitochondrial pseudogenes and common SNPs were downloaded from the UCSC table browser. CAGE tags in A549 cells, generated by the ENCODE project, were obtained from http://ccg.vital-it.ch/mga/hg19/encode/GSE34448/A549_cell_longPolyA_rep1.sga.gz. Intersections between genomic coordinates were performed using bedtools(Quinlan, 2014).

To assess the stop rates across the various classes of detected sites, we utilized the m1A-seq-SuperScript datasets generated in (Safra et al., 2017). The stop rate is calculated as the fraction of reads beginning at each site, divided by the number of reads overlapping it (Safra et al., 2017).

To visually inspect and reanalyze the Li et al data, raw data was downloaded from GEO: GSE102040. The reads from Input, IP, and IP+demethylase samples (2 replicates each) were aligned to the human genome (assembly: hg19) using STAR aligner (Dobin et al., 2013). To clip the 10 nt barcode at the 5’ end, and the adapter at the 3’ end, we used the following criteria ‘--clip3pAdapterSeq AGATCGGAAGAGCGTCGTGTAGGGAAAGAGTGT --clip5pNbases 10’. The datasets were analyzed using the approach described in (Safra et al., 2017).

The Venn-diagrams comparing the putative m1A sites with previously defined peaks were done on the basis of an intersection of the transcriptome-level coordinates provided for the ‘m1A peaks’, downloaded as a Supplementary Table from (Li et al., 2016). For the comparison with the Dominissini et al paper, we defined a window of 200 nt centered around the center of the 2129 ‘m1A peaks’ which were reported for HEK293 cells, downloaded from the Supplementary Tables in (Dominissini et al., 2016).

## References

Bazak, L., Haviv, A., Barak, M., Jacob-Hirsch, J., Deng, P., Zhang, R., Isaacs, F.J., Rechavi, G., Li, J.B., Eisenberg, E., et al. (2014). A-to-I RNA editing occurs at over a hundred million genomic sites, located in a majority of human genes. Genome Res. 24, 365–376.

Carlile, T.M., Rojas-Duran, M.F., Zinshteyn, B., Shin, H., Bartoli, K.M., and Gilbert, W.V. (2014). Pseudouridine profiling reveals regulated mRNA pseudouridylation in yeast and human cells. Nature 515, 143–146.

Carninci, P., Sandelin, A., Lenhard, B., Katayama, S., Shimokawa, K., Ponjavic, J., Semple, C.A., Taylor, M.S., Engstrom, P.G., Frith, M.C., et al. (2006). Genome-wide analysis of mammalian promoter architecture and evolution. Nat. Genet. 38, 626–635.

Chang, H., Lim, J., Ha, M., and Kim, V.N. (2014). TAIL-seq: genome-wide determination of poly(A) tail length and 3' end modifications. Mol. Cell 53, 1044–1052.

Chen, D., and Patton, J.T. (2001). Reverse transcriptase adds nontemplated nucleotides to cDNAs during 5'-RACE and primer extension. Biotechniques 30, 574–580, 582.

Dobin, A., Davis, C.A., Schlesinger, F., Drenkow, J., Zaleski, C., Jha, S., Batut, P., Chaisson, M., and Gingeras, T.R. (2013). STAR: ultrafast universal RNA-seq aligner. Bioinformatics 29, 15–21.

Dominissini, D., Moshitch-Moshkovitz, S., Schwartz, S., Salmon-Divon, M., Ungar, L., Osenberg, S., Cesarkas, K., Jacob-Hirsch, J., Amariglio, N., Kupiec, M., et al. (2012). Topology of the human and mouse m6A RNA methylomes revealed by m6A-seq. Nature 485, 201–206.

Dominissini, D., Nachtergaele, S., Moshitch-Moshkovitz, S., Peer, E., Kol, N., Ben-Haim, M.S., Dai, Q., Di Segni, A., Salmon-Divon, M., Clark, W.C., et al. (2016). The dynamic N(1)-methyladenosine methylome in eukaryotic messenger RNA. Nature.

Hauenschild, R., Tserovski, L., Schmid, K., Thüring, K., Winz, M.-L., Sharma, S., Entian, K.-D., Wacheul, L., Lafontaine, D.L.J., Anderson, J., et al. (2015). The reverse transcription signature of N-1-methyladenosine in RNA-Seq is sequence dependent. Nucleic Acids Res. 43, 9950–9964.

Haussmann, I.U., Bodi, Z., Sanchez-Moran, E., Mongan, N.P., Archer, N., Fray, R.G., and Soller, M. (2016). m6A potentiates Sxl alternative pre-mRNA splicing for robust Drosophila sex determination. Nature.

Ledergerber, C., and Dessimoz, C. (2011). Base-calling for next-generation sequencing platforms. Brief. Bioinform. 12, 489–497.

Lence, T., Akhtar, J., Bayer, M., Schmid, K., Spindler, L., Ho, C.H., Kreim, N., Andrade-Navarro, M.A., Poeck, B., Helm, M., et al. (2016). m6A modulates neuronal functions and sex determination in Drosophila. Nature.

Li, X., Zhu, P., Ma, S., Song, J., Bai, J., Sun, F., and Yi, C. (2015). Chemical pulldown reveals dynamic pseudouridylation of the mammalian transcriptome. Nat. Chem. Biol. 11, 592–597.

Li, X., Xiong, X., Wang, K., Wang, L., Shu, X., Ma, S., and Yi, C. (2016). Transcriptome-wide mapping reveals reversible and dynamic N1-methyladenosine methylome. Nat. Chem. Biol.

Li, X., Xiong, X., Zhang, M., Wang, K., Chen, Y., Zhou, J., Mao, Y., Lv, J., Yi, D., Chen, X.-W., et al. (2017). Base-Resolution Mapping Reveals Distinct m1A Methylome in Nuclear- and Mitochondrial-Encoded Transcripts. Mol. Cell 68, 993–1005.e9.

Meyer, K.D., Saletore, Y., Zumbo, P., Elemento, O., Mason, C.E., and Jaffrey, S.R. (2012). Comprehensive Analysis of mRNA Methylation Reveals Enrichment in 3' UTRs and near Stop Codons. Cell 149, 1635–1646.

Meyer, K.D., Patil, D.P., Zhou, J., Zinoviev, A., Skabkin, M.A., Elemento, O., Pestova, T.V., Qian, S.-B., and Jaffrey, S.R. (2015). 5' UTR m 6 A Promotes Cap-Independent Translation. Cell 163, 999–1010.

Plessy, C., Bertin, N., Takahashi, H., Simone, R., Salimullah, M., Lassmann, T., Vitezic, M., Severin, J., Olivarius, S., Lazarevic, D., et al. (2010). Linking promoters to functional transcripts in small samples with nanoCAGE and CAGEscan. Nat. Methods 7, 528–534.

Quinlan, A.R. (2014). BEDTools: the Swiss-army tool for genome feature analysis. Curr. Protoc. Bioinformatics 11–12.

Safra, M., Sas-Chen, A., Nir, R., Winkler, R., Nachshon, A., Bar-Yaacov, D., Erlacher, M., Rossmanith, W., Stern-Ginossar, N., and Schwartz, S. (2017). The m(1)A landscape on cytosolic and mitochondrial mRNA at single-base resolution. Nature.

Schwartz, S., Agarwala, S.D., Mumbach, M.R., Jovanovic, M., Mertins, P., Shishkin, A., Tabach, Y., Mikkelsen, T.S., Satija, R., Ruvkun, G., et al. (2013). High-resolution mapping reveals a conserved, widespread, dynamic mRNA methylation program in yeast meiosis. Cell 155, 1409–1421.

Schwartz, S., Bernstein, D.A., Mumbach, M.R., Jovanovic, M., Herbst, R.H., León-Ricardo, B.X., Engreitz, J.M., Guttman, M., Satija, R., Lander, E.S., et al. (2014a). Transcriptome-wide mapping reveals widespread dynamic-regulated pseudouridylation of ncRNA and mRNA. Cell 159, 148–162.

Schwartz, S., Mumbach, M.R., Jovanovic, M., Wang, T., Maciag, K., Bushkin, G.G., Mertins, P., Ter-Ovanesyan, D., Habib, N., Cacchiarelli, D., et al. (2014b). Perturbation of m6A writers reveals two distinct classes of mRNA methylation at internal and 5' sites. Cell Rep. 8, 284–296.

Shi, H., Wang, X., Lu, Z., Zhao, B.S., Ma, H., Hsu, P.J., and He, C. (2017). YTHDF3 facilitates translation and decay of N(6)-methyladenosine-modified RNA. Cell Res.

Takahashi, H., Kato, S., Murata, M., and Carninci, P. (2012). CAGE (cap analysis of gene expression): a protocol for the detection of promoter and transcriptional networks. Methods Mol. Biol. 786, 181–200.

Wang, X., Lu, Z., Gomez, A., Hon, G.C., Yue, Y., Han, D., Fu, Y., Parisien, M., Dai, Q., Jia, G., et al. (2014). N6-methyladenosine-dependent regulation of messenger RNA stability. Nature 505, 117–120.

Wang, X., Zhao, B.S., Roundtree, I.A., Lu, Z., Han, D., Ma, H., Weng, X., Chen, K., Shi, H., and He, C. (2015). N(6)-methyladenosine Modulates Messenger RNA Translation Efficiency. Cell 161, 1388–1399.

Whiteford, N., Skelly, T., Curtis, C., Ritchie, M.E., Löhr, A., Zaranek, A.W., Abnizova, I., and Brown, C. (2009). Swift: primary data analysis for the Illumina Solexa sequencing platform. Bioinformatics 25, 2194–2199.

Zhou, J., Wan, J., Gao, X., Zhang, X., Jaffrey, S.R., and Qian, S.-B. (2015). Dynamic m6A mRNA methylation directs translational control of heat shock response. Nature 526, 591–594.

